# Explainable Graph Learning for Multimodal Single-Cell Data Integration

**DOI:** 10.1101/2024.12.06.627151

**Authors:** Mehmet Burak Koca, Fatih Erdoğan Sevilgen

## Abstract

Integrating multi-omic single-cell data is essential for uncovering cellular heterogeneity and identifying specialized subpopulations. However, achieving both explainable and expressive integration remains challenging due to the complex relationships between modalities. Here, we introduce Single-Cell PROteomics Vertical Integration (SCPRO-VI), a novel algorithm that integrates paired multi-omic data through similarity graph fusion, enhanced with a multi-view variational graph auto-encoder. SCPRO-VI incorporates a biologically guided distance metric and a multi-view graph-based embedding approach to effectively capture cross-modality relations. Extensive benchmark on multi-omic CITE-seq datasets shows that SCPRO-VI significantly enhances inter-cell type heterogeneity and identifies biologically meaningful sub-clusters that remain indistinguishable by existing methods. These results demonstrate robustness of SCPRO-VI and its potential to address key challenges in single-cell multi-omic data integration.

## 1 Introduction

The regulatory dynamics among biological molecules across different omic layers remain far from fully understood. These complex interactions are often non-linear and vary significantly across cell types [1]. Recent advancements in single-cell technologies have enabled the simultaneous (paired) sequencing of multiple omic layers within the same cells [2], offering deeper insights into cellular complexity. Integrating multi-omic data facilitates the identification of cell-type-specific correlations among these molecules, and conversely, these patterns can be leveraged to distinguish between different cell types [3]. Furthermore, such integrated analyses can reveal specialized sub-clusters within the same cell type, providing a higher resolution of cellular heterogeneity.

Integrating cell views from different modalities (vertical) presents unique challenges compared to the more established uni-modal (horizontal) integration of single cells across different datasets [4]. In multi-modal data integration, the primary goal is to establish optimal alignment between the naturally varied modality views of cells, whereas horizontal integration focuses on minimizing technical variations between uni-modal datasets. Constructing a unified representation from multi-omic data is particularly challenging, as each modality often introduces distinct relationships between cells, making it difficult to ascertain which associations are most accurate for each cell type [3]. These challenges are further amplified when the modalities are not paired [5]. Several methods have been developed to address the single-cell multi-omic data integration problem [6]. These approaches are categorized into statistical modeling [7], similarity graph fusion [3, 8], and embedding into a joint latent space [1, 5, 9–16].

Statistical modeling methods integrate multi-omic data by fitting individual omic distributions based on assumed correlations. However, their reliance on predefined assumptions often limits their ability to capture the complexity and variability of biological systems.

Similarity graph fusion methods calculate omic-specific similarities among cells for each modality individually and then combine these into a unified similarity graph [3, 8]. These approaches are valuable for analyzing agreements and conflicts between omic layers and allow for assessing cell-type-specific contributions of modalities by comparing fused similarities to modality-specific graphs. However, fusing similarity graphs is not trivial. These methods must account for differences in scale and distribution of similarities to prevent one modality from dominating the fusion due to numerical imbalances. Additionally, the generation of joint cell embeddings must carefully preserve fused similarities to avoid significant information loss in downstream analyses.

Joint embeddings of modalities into a shared latent space are achieved using non-negative matrix factorization (NMF) [9–11], canonical correlation analysis (CCA) [12], and deep learning methods [1, 5, 13–16]. NMF techniques extract low-dimensional representations that identify shared and dataset-specific factors across single-cell multi-omics datasets. While NMF is useful for interpreting modality contributions, its linear nature makes it less effective in capturing non-linear relationships. CCA integrates datasets by identifying linear combinations of variables that maximize correlations and explain variability within and between datasets. Although effective for paired datasets, CCA’s performance is often constrained by variability in modality distributions.

Deep learning methods have emerged as powerful tools for integrating modalities into a shared latent space [17]. These approaches employ various architectures such as auto-encoders [1, 13, 14], convolutional neural networks [15], generative adversarial networks [5], and graph neural networks [16]. By capturing non-linear connections between features, deep learning overcomes the limitations of linear methods, making it particularly suited for multi-omic data integration. However, a significant challenge lies in the limited explainability and interpretability of the resulting joint embeddings, which provide poor insight into inter-modality relationships.

Here, we present the Single-Cell PROteomics Vertical Integration (SCPRO-VI) method, a similarity graph fusion approach that incorporates a multi-view variational graph auto-encoder (VGAE) for embedding modalities into a latent space. SCPRO- VI integrates a novel metric that includes biological guidance into the construction of similarity graphs. These modality-specific similarity graphs are fused into a unified similarity graph by normalizing and balancing the local neighborhoods of cells across modalities. The resulting unified graph is then used to generate integrated cell embeddings through a novel multi-view VGAE model, ensuring that the embeddings accurately capture cell similarities as represented in the fused graph. Thus, SCPRO-VI stands as a powerful data integration tool that leverages the capabilities of deep learning for integration while providing explainable and interpretable relationships among cells that shape the integrated space.

SCPRO-VI was rigorously evaluated on a multi-omic CITE-seq dataset [3] to assess its efficiency and effectiveness. Additionally, we benchmarked our algorithm against state-of-the-art single-cell multi-omic data integration tools [3, 9, 11, 13] in extensive experiments using a challenging multi-omic CITE-seq dataset [20] characterized by low inter-cell type heterogeneity. While our experiments focused on integrating proteomics and transcriptomics data available in these datasets, it is important to note that SCPRO-VI is versatile and can integrate any paired modalities with numerical features, with or without biological guidance. The experimental results show that SCPRO-VI enhances inter-cell type heterogeneity more effectively than competing methods, as supported by quantitative performance metrics. Furthermore, in-depth analyzes of the integrated cell embeddings revealed that SCPRO-VI effectively identifies specialized cell subgroups that remain indistinguishable in the integration results of other methods.

## 2 Results

### 2.1 Integrating paired multi-omics in single-cells using variational graph auto-encoders

Single-Cell PROteomics Vertical Integration (SCPRO-VI) is a computational tool for single-cell multi-omic data integration, specifically tailored for integrating proteomic data with transcriptomic data.

SCPRO-VI introduces a biologically reinforced distance metric for calculating distance between two cells and a novel multi-view variational graph auto-encoder (VGAE) for integrating modalities as cell embeddings by using the novel distances. The distance metric measures the alignment of complete graphs across modality features for each cell. The alignment score is the sum of differences between weights of the same edges(see Methods). Since every relation between features isn’t in the same importance [18], our method enables weighting each edge alignment by a biological relevance score (see Methods). Here, we used protein-protein interaction scores obtained from the STRING [19] database for weighting the relations between protein pairs while calculating cell-to-cell distance matrix *D^P^* by proteomics. Thus, our metric reflects how two cells are distant by their protein-protein interaction networks on cell membranes which is expected to be low between the same type of cells [3].

Supervised biological guidance beyond the vectorial similarity of cell features enhances the heterogeneity among the cells by their cell types. The distance method reduces the impact of less critical proteins for cell type differentiation. Moreover, highlighting active proteins enhances clustering accuracy among diverse cell types (see Supplementary Fig. 1A). We employed gene ontology (GO) analysis for the proteins measured in our test datasets [3, 20]. For each dataset, the most and least important protein sets are identified, and GO analysis was applied for each of them individually (see Methods). The analysis results showed that the proteins with higher interaction scores are more active in distinctive biological processes than the proteins appeared in weak interactions (see Supplementary Fig. 1B).

The important proteins in the HIV dataset [3] were annotated as playing a more active role in the immune system (*P* = 1.084∗10^−32^) than the least important proteins (*P* = 8.445 ∗ 10^−17^). Moreover, the important proteins were annotated with GO:MF virus receptor term by a higher score (*P* = 7.861 ∗ 10^−20^) than the least important proteins (*P* = 8.303 ∗ 10^−6^). All these results suggest weighting the proteins while calculating the distances between cells potentially increases the performance of matching the same type of cells conducting the same biological processes.

The biological guidance should be determined carefully since a ranking for the relevance of features isn’t available for all modalities. In our experiments, we tested the number of co-occurrences in pathways as relevance scores for gene pairs to calculate transcriptomic-based distance matrix *D^T^* . However, using such a relevance score caused a negative effect on inter-cell type heterogeneity (see Supplementary Fig. 1C). Thus, we decided not to use the metric for transcriptomics, instead we applied dimensionality reduction to raw transcriptomics with Principle Component Analysis (PCA). Then, *D^T^* is calculated by the cosine distance between PCA components of cell pairs (see Methods).

The distance matrices are normalized to prevent over-considering a modality. Each cell’s distances are normalized by the mean of its k-nearest neighbors (see Methods). The value of *k* limits the size of neighborhood considered while integrating modalities. The normalized distance matrices are element-wise multiplicated to have an integrated distance matrix *D^I^* (Fig. 1A).

**Fig. 1.**
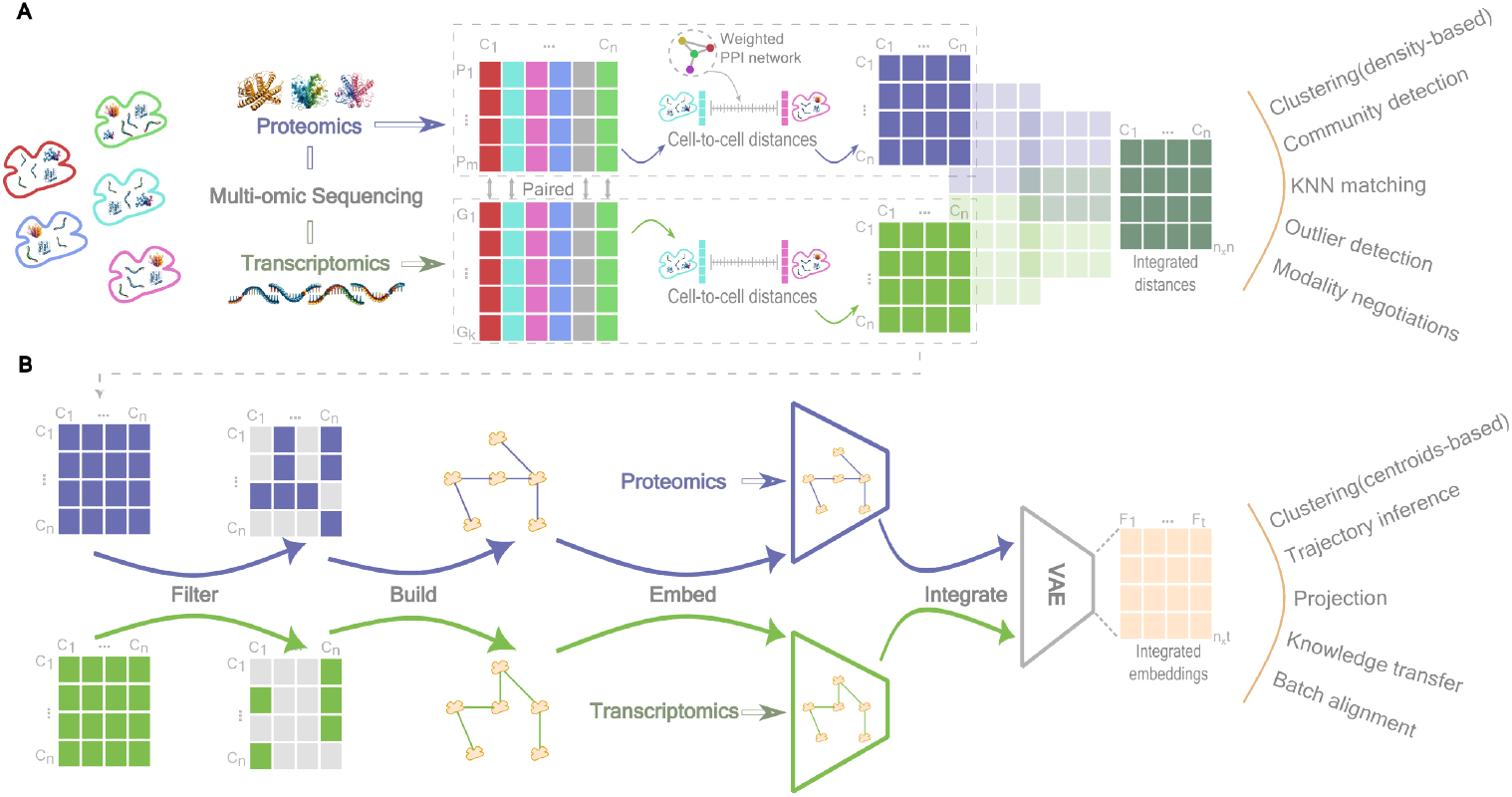
A. Schematic representation of modality-based distance matrix calculation and integrated distances generation; **B** Schematic representation of generating integrated cell embeddings using multi-view variational graph auto-encoders.

The cell distances can facilitate community detection, k-nearest neighbor (k-nn) analysis, outlier detection, evaluation of modality contributions to cell differentiation, and analysis of modality importance for each cell type [3]. Density-based clustering can also be applied using only cell distances, as these algorithms rely on neighborhood relationships [21]. However, embedding this distance information into a latent space is essential for downstream tasks that require vectorial cell representations, such as centroid-based clustering or knowledge transfer [1].

SCPRO-VI introduce a novel variational graph auto-encoder (VGAE) model (see Methods) for generating integrated cell embeddings (Fig. 1B). We employ graph auto- encoders as they are developed primarily to encode neighborhood information into the embedding space, aligning with our integration goals. The model employs a multi-view approach, using separated GVAE models for each modality while sharing the same vertex set *G* (all cells in the dataset) with modality-specific edge sets *E^P^* and *E^T^* . The edge sets *E^P^* and *E^T^* are generated by thresholding the distances *D^P^* and *D^T^*, respectively (see Methods). Raw proteomics and PCA embeddings of transcriptomics measurements are used as node features within their respective VGAE models. The models are individually pre-trained to achieve faster and more stable convergence when generating integrated embeddings. Using variational inference instead of traditional auto-encoders improves integration consistency, as it facilitates combining similar distributions rather than unrelated latent spaces [22].

The modality-specific cell embeddings from pre-trained models are fused by an auto-encoder model. The auto-encoder has no decoder part; instead, the pairwise distances between latent cell embeddings are calculated to determine loss (see Methods). The cumulative difference between element-wise distances in the predicted distances and *D^I^*is computed as the loss (see Methods) and all parameters including GVAEs are updated accordingly. Thus, the end-to-end model is trained to generate integrated embeddings that reflect the pairwise relations in the integrated distances.

These embeddings allow for transfering the multi-omic neighborhood perspective to the downstream tasks [13, 23].

The effectiveness and efficiency of SCPRO-VI were evaluated through extensive experiments. We first tested our algorithm’s robustness on a comprehensive dataset containing 24 samples from 8 patients [3]. State-of-the-art multi-omics data integration algorithms Seurat v4 [3], and Mofa+ [9] were also included in tests for a relative assessment of our algorithm’s efficiency. SCPRO-VI was further benchmarked against state-of-the-art algorithms TotalVI [13], Mowgli [11], Seurat v4 [3], and Mofa+ [9] on a more complex dataset with fewer cell type heterogeneity. Moreover, several bioinformatics analyses were employed to prove the effectiveness of our integration algorithm. Although many other state-of-the-art algorithms exist, we selected widely used algorithms in all types that offer readily accessible code packages for testing.

### 2.2 Validation of SCPRO-VI robustness over systematically generated multi-sample benchmark datasets

The performance of the SCPRO-VI was evaluated on a dataset [3] generated by combining cells sequenced in 24 individual experiments to assess its reusability across different datasets. The dataset was generated by the authors of the Seurat v4 [3] algorithm, gathering samples from 8 patients with HIV infection. The samples were obtained from each volunteer at three time points: the day before vaccination, and three and seven days post-vaccination. The samples were frozen and sequenced at once to avoid batch effect as much as possible. The potential differentiation between time points and the variations among volunteers enables assessment of our algorithm’s resilience to biological variations without concerning the technical variations or noise. We measured the performance of the integrated embeddings using five different widely-used metrics in data integration studies: 1-NN cell matching, silhouette score, adjusted rand index (ARI), normalized mutual information (NMI), and graph connectivity (see Methods). The state-of-the-art benchmark algorithms Seurat v4 and Mofa+ were also tested and evaluated to assess our performance results comparatively. The performance metrics were also applied to the concatenated transcriptomics and proteomics features of raw data to establish a baseline for performance metrics. TotalVI algorithm couldn’t be tested since it requires unnormalized protein counts which is not available in this dataset. Another benchmark algorithm Mowgli also excluded since it didn’t show promising results in other experiments despite its long execution times.

Moreover, comprehensive comparisons were employed to benchmark our algorithm against state-of-the-art algorithms, which is not the case in this experiment.

Algorithm test results are presented as a heat map to facilitate tracking robustness across time points and between samples (Fig. 2). In addition, the mean performances of algorithms for each time point are given in the heat map for an overview of vaccination effect. Our algorithm results mostly shaded in green tones indicating high performance. Seurat algorithm also obtained greeny results in all metrics except for the ARI metric. However, the Mofa+ algorithm performed poorly across all metrics except graph connectivity.

**Fig. 2.**
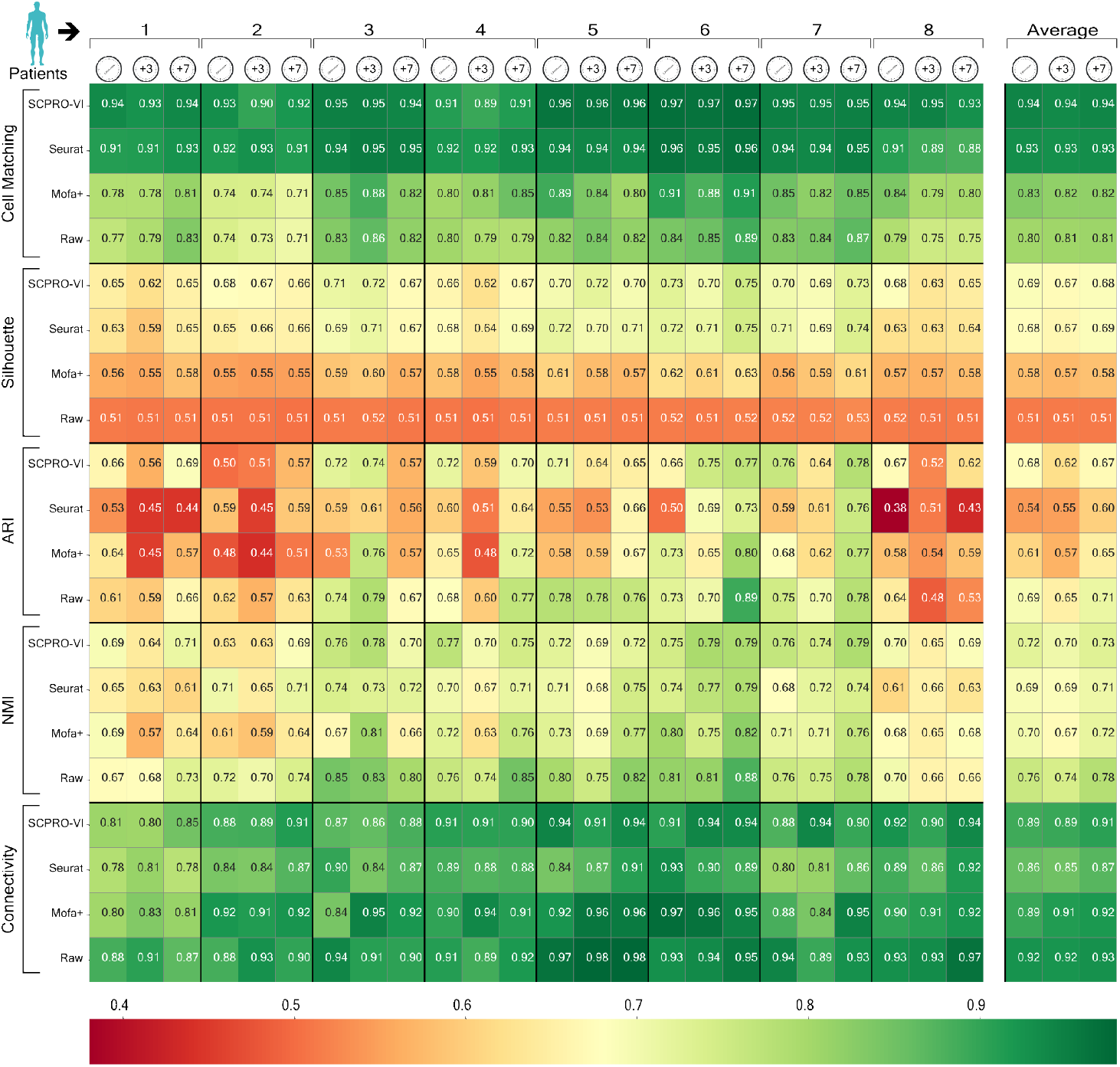
Performance results of integration algorithms in five metrics as heat map. Each column contains results for a sample. Every three consecutive columns belong to a single patient. The samples of patients are ordered as before vaccination, 3 days after vaccination, and 7 days after vaccination. Rows represent algorithms and are grouped by the metrics.

All three algorithms improve the baseline at cell matching and silhouette score metrics. The cell matching score proved that our algorithm has consistent success at gathering the cells of the same type together by 0.3% variation at most between the matching performance of three samples of a patient. SCPRO-VI significantly improved baseline cell matching, achieving a 9% to 21% performance increase. Seurat also shows similar performance for gathering the same cells together, but still, our algorithm performs 1% better than Seurat at mean performance and the performance improvement rises 5% for the patient 8. Mofa+ improves the performance of baseline up to 7% but mostly has the same performance with raw data.

Silhouette scores indicate that SCPRO-VI enhances inter-cluster heterogeneity across all patients. These significant improvements over baseline rise to 23%. Seurat algorithm has a similar performance result as in the cell matching since their algorithm tends to separate cells in clusters with less concerning intra-cluster fragmentations.

Although Mofa+ improved raw data silhouette scores, it remained inferior to SCPRO- VI and Seurat. ARI and NMI metrics indicate that none of the algorithms achieved clustering more aligned with ground-truth cell clustering than the raw data. However, SCPRO-VI has 2% better NMI scores when compared with both Seurat and Mofa+ where they have similar performances. On the other hand, SCPRO-VI outperforms both algorithms in distinguishing cells that should not cluster together, as reflected in the ARI metric, unlike NMI (see Methods). Our algorithm has 9% and 5% better ARI scores at mean from Seurat and Mofa+, respectively.

Graph connectivity metric should be evaluated by considering other performance metrics since it isn’t sensitive to joint clustering of different cell types (see Methods). The results indicate that the same type of cells are close together in raw data, but they are mostly intersecting cell clusters of different types. On the other hand, the integration algorithms strive to separate different types of cell clusters for increasing heterogeneity. Therefore, the graph connectivity scores of both SCPRO-VI and Mofa+ are similarly 2% lower than raw data. The Seurat algorithm had the worst results with a 6% lower score than raw data since it tends to generate more than one sub-cluster for a cell type.

Our algorithm obtained robust and relatively higher performance results in all the performance metrics. Overall, the results show that SCPRO-VI balances reducing intra-cluster fragmentation with enhancing inter-cluster heterogeneity. In addition, the algorithm showed its stability by having the lowest standard deviation between results in all metrics. The findings suggest that SCPRO-VI is robust and well-suited for vertical integration of the multi-omic datasets obtained from diverse samples across different biological conditions.

### 2.3 Multi-omic inference of variations at HIV patients pre- and post-vaccination

Transcriptomics and proteomics expression profiles are expected to change after vaccination [3]. Studies report upregulation or downregulation of certain genes and proteins after vaccination in HIV patients [24]. These genes and proteins shape specialized cell sub-clusters set forth on various tasks in the immune system [25].

We examined our integration results to identify cell subsets emerging post-vaccination. This approach enabled us to test our algorithm’s effectiveness in differentiating novel cell sub-clusters, one of the primary goals of vertical integration. The UMAP projections show that several sub-clusters appeared after vaccination for all patients (see Supplementary Fig. 2). There were both immediately emerging clusters on day 7 and gradually separating clusters that began to diverge on day 3 were fully revealed by day 7.

We select a gradually separating monocyte sub-cluster for further analysis (Fig. 3A). While several sub-clusters were available, the analyzed cluster is arbitrarily selected as it visually distinct. Additionally, monocytes are well-studied in HIV infection, facilitating interpretation of potential changes in expression levels. First, we clustered cells to identify those within the targeted sub-cluster (see Methods). Next, we identified the most distinctive proteins and genes contributing to intra-cell-type separation (see Methods and Supplementary Fig. 3A).

**Fig. 3.**
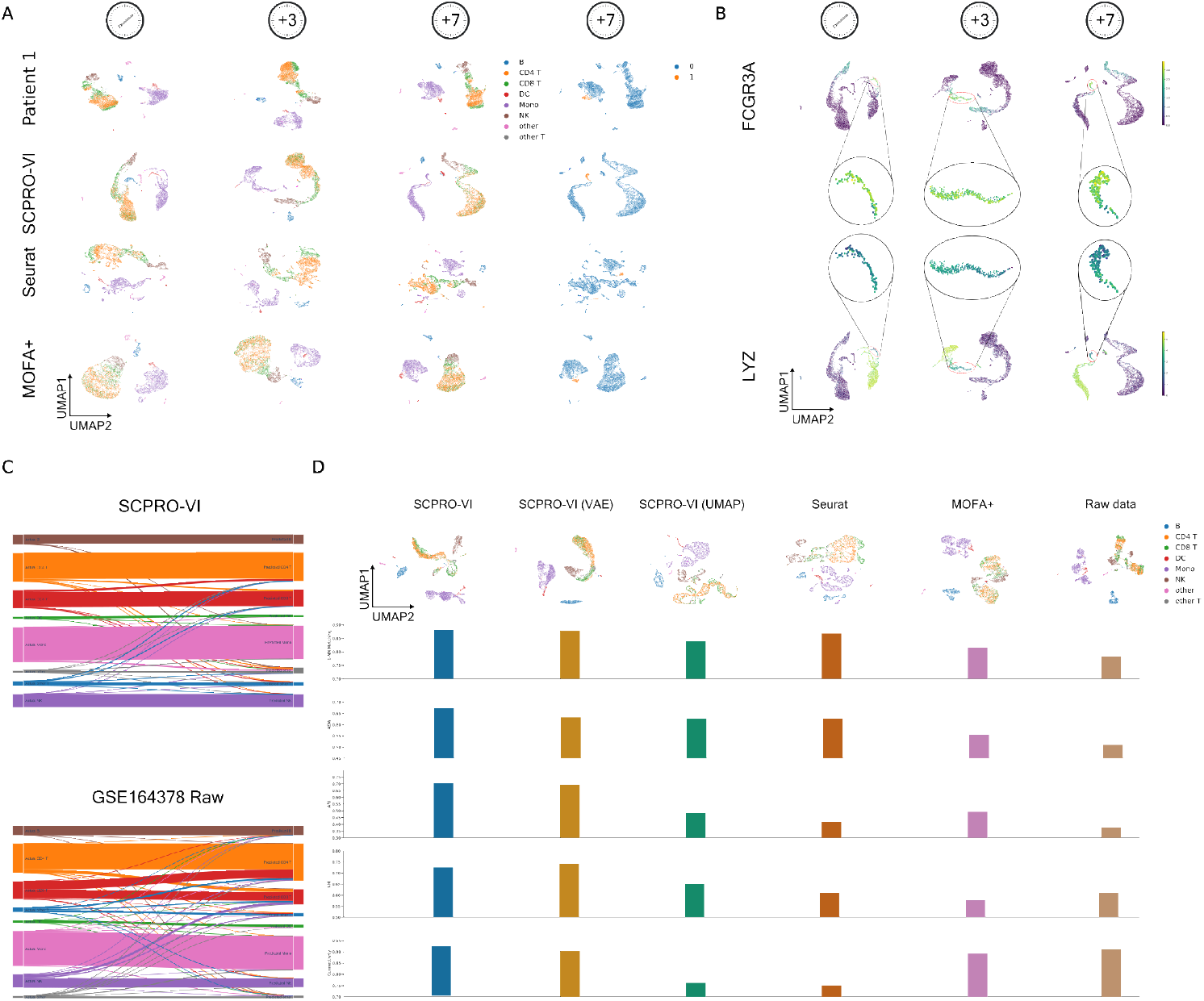
The effectiveness test results of our integration algorithm on the HIV dataset. **A** UMAP projections of patient 1 raw and integrated measurements for the datasets sequenced before vaccination, 3 days after, and 7 days after vaccination. The analyzed monocyte cell sub-cluster is colored on raw data and algorithm embeddings for the 7th-day dataset (right). **B** Umap projections of SCPRO-VI embeddings for all datasets of patient 1 which are colored by expression levels of FCGR3A protein (top) and LYZ gene (bottom). **C** The Sankey diagrams of matching actual type of cells (left) with their nearest neighbors (right) by both SCPRO-VI embeddings (top) and raw data in the whole HIV dataset (bottom). **D** The UMAP projections (top) and numerical performance results (bottom) for the integration of the sub-sampled HIV dataset.

In the feature analysis, FCGR3A was found as the most important protein at the separation of the sub-cluster from the rest monocytes. This protein is a well-known marker playing an active role in HIV transmission and infection processes [26, 27]. However, when the analysis was repeated for monocytes versus all cells, this protein ranked 80th in importance among 174 proteins (see Methods). Therefore, we concluded that these monocyte cells may be specialized for a specific function related to FCGR3A.

We also analyzed genes and identified LYZ as the most distinctive gene for this sub- cluster. This gene was reported as one of five key markers for differentiating patients who respond positively to HIV treatment from those with negative responses [28].

Furthermore, LYZ was ranked as the fifth most important gene for distinguishing monocytes from other cells, making it challenging to identify this specialized sub- cluster based solely on gene expression.

We examined the expression levels of the FCGR3A protein and LYZ gene across all cells to analyze their expression profiles in various cell types (Fig. 3B). We observed that FCGR3A is upregulated exclusively in the target sub-cluster and downregulated in all natural killer (NK) cells. Conversely, the LYZ gene is active only in monocytes and dendritic cells (DC), with upregulation in both but specific downregulation in cells within the analyzed sub-cluster. These results indicate that the co-occurrence of FCGR3A upregulation and LYZ downregulation is unique to the target sub-cluster.

Both distinctive feature analyses and expression profile analyses support that the target cell sub-cluster reveals by integrating transcriptomics and proteomics data. Additionally, the LYZ gene and FCGR3A protein have been identified as co-differentiating markers of CD16+ monocyte sub-clusters in studies on immune cell exhaustion in HIV-infected patients [25]. Together, this evidence highlights SCPRO- VI’s effectiveness in multi-omic integration for identifying specialized cell sub-clusters that emerge uniquely through multi-omic data integration.

We also reviewed the raw concatenated data of patient 1 and the integration results from other algorithms to examine similar gradual cell grouping (see Fig. 3A). The raw data showed no significant changes in cell grouping across the three time points. However, cells within the analyzed sub-cluster appeared closer in the day 7 measurements, though they did not form a distinct sub-cluster. Mofa+ generated gradually narrowing cell clusters, but the target cells did not fully separate from other monocytes, despite drawing closer. Seurat generated multiple monocyte sub-clusters in all three datasets. However, the projection of the target sub-cluster showed that the algorithm divided it into two distinct clusters. These findings suggest that revealing specialized sub-clusters through multi-omic integration is challenging, yet achievable with our algorithm.

### 2.4 Vertical integration of modalities in cells from multi-samples

Integrating cells from multiple samples can reduce performance due to biological variations that introduce mixed cell populations. We evaluated our algorithm’s performance on the complete multi-sample dataset by calculating 1-NN cell matchings based on cell type using integrated distances (Fig. 3C). We compared SCPRO-VI’s matching results with concatenated raw data to assess improvements in inter-cell heterogeneity. We used the cosine distance to calculate cell-to-cell distances, as it produced the best results among the common distance metrics (see Supplementary Fig. 3B).

The Sankey diagram shows the actual type of cells (left) and their 1-NN neighbor matching (right). The diagram overview highlights the decrease in mismatches, shown by fewer crossing lines. The number of mismatched cells by the raw data is significantly reduced in 5 out of 8 cell types with our integrated distances (see Supplementary Fig. 3C). Our algorithm slightly reduced matching accuracy by 0.025% for B cells, 0.05% for DC cells, and 0.004% for monocytes. The mismatches in DC and monocyte cells are likely due to errors in uncertain regions, such as cluster edges. However, almost all (413 of the 425) mismatches for B cells occurred with Other T cells. We observed that the mismatched cells are close to the Other T cells, separated from the main B cells cluster (see Supplementary Fig. 3D). Our algorithm struggled to differentiate these intersected B cells from Other T cells, as they closely resemble Other T cells despite being labeled as B cells.

Raw data mismatched 6066 out of 18664 NK cells. The most mismatched targets were in CD8 T cell type by 3366 mismatches. SCPRO-VI reduced the number of mismatches to 371. The total matching performance was significantly increased from 67% to 98%, which makes it much easier the distinguish them from the T cells. Lastly, Other T cells mismatched 35% with CD4 T cells and 23% with CD8 T cells in the raw data. Our algorithm achieved to reduce the mismatch rate of Other T cells with CD4 T cells to 8% and with CD8 T cells to 7%.

Our algorithm achieved the highest improvements at Other T and CD8 T cell types by 88% and 70%, respectively. CD8 T cells had a 52% mismatch rate with other cell types. The CD8 T cells were intersected with CD4 T cells in raw data resulting in 92% of CD8 T mismatches being established with CD4 T cells. The proposed distance metric reduced mismatches between CD8 and CD4 T cells by 88%, allowing clear differentiation of CD8 T cells.

We conducted an additional experiment to assess our metric’s contribution to CD8 T cell matching accuracy. We calculated integrated distances using cosine distances for both modalities instead of calculating proteomic distances with our distance metric. The results showed that the mismatching ratio of CD8 T cells increased from 17% to 25%. These findings suggest that our novel distance metric enhances cell matching accuracy. Moreover, results further indicated that integration was more effective when modalities were handled individually rather than concatenated. Together, these results demonstrate our algorithm’s effectiveness in clustering similar cell types, raising the overall cell matching accuracy from 80.7% to 91.3%.

The integration performance of SCPRO-VI on multi-sample datasets was also measured using additional performance metrics. However, we randomly sampled 5k cells from the 161764 cells in the HIV dataset to maintain a consistent size with the individual samples (see Methods). The Seurat and Mofa+ algorithms were included as in the previous test on this dataset (Fig. 3D). Moreover, we tested a variational auto-encoder (VAE) version of our model that retains the same architecture, but linear layers were used instead of graph convolution layers (see Methods). Lastly, we tested UMAP embeddings generated using integrated distances (see Methods). The VAE model and UMAP embeddings were included in the comparison for assessing the contribution of using a graph auto-encoder model.

SCPRO-VI achieved 1% increase in cell matching accuracy compared to the linear model, and a 4% increase over UMAP embeddings. Furthermore, SCPRO-VI outperformed the Seurat algorithm, achieving a matching accuracy of 88.1% which is 1.5% higher than that achieved by Seurat. Mofa+ managed to match the cells with an accuracy of 81.5%, improving the raw data performance by only 2%.

SCPRO-VI also demonstrated the best performance in terms of the average silhouette score and the Adjusted Rand Index (ARI), with improvements of 5% and 1%, respectively, compared to the linear model. However, the linear model outperformed by 1.5% in NMI and by 2% in graph connectivity metrics. These results suggest that the linear model generates embeddings with smaller intra-cluster distances, as indicated by both numerical results and UMAP projections. In contrast, the silhouette and ARI scores indicate that the inter-cell heterogeneity is reduced in the linear model compared to SCPRO-VI. This observation aligns with the inherent differences between the two models, as the Variational Graph Autoencoder (VGAE) model differentiates unrelated cells through both its loss function and the aggregation process used in convolution.

Comparison results further illustrate the superiority of deep learning models for embedding integrated distances over UMAP. Additionally, these results suggest that integration performance improves when the model is informed by neighborhood information, which helps to minimize unwanted similarities between unrelated cell pairs.

### 2.5 Benchmarking against state-of-art algorithms on a complex dataset with weak inter-cluster heterogeneity

The multi-omic integration capabilities of the SCPRO-VI algorithm were compared against state-of-the-art algorithms using a complex paired multi-omic dataset [20]. We chose this dataset for our benchmark due to the poor separation between different types of cell clusters, which presents a challenge for the enhancing inter-cell heterogeneity goal of integration.

The benchmark dataset [20] comprises 28k mononuclear cells (MNCs) in 12 cell types sampled from 14 individuals. Bone marrow samples were obtained from four adults and four fetuses across different developmental stages. Liver samples were collected from six fetuses. Additionally, four samples were obtained from newborn cord blood. The multi-sample and multi-organ structure of this dataset resulted in separating the same type of cells into more than one cluster. However, the clusters exhibit substantial overlap, since they are all MNCs. These characteristics make the dataset a suitable candidate for benchmarking multi-omic data integration.

We compared our integration algorithm with Seurat v4 [3], Mofa+ [9], and the concatenation of raw data as a baseline, using the same setup used for the MNC dataset integration. Seurat is the most widely used benchmark algorithm in multi-omic data integration studies. This method focuses on the integration of neighborhoods across different modalities, aligning well with our integrated distance approach, making it a strong candidate for comparison. Mofa+ is another powerful tool for vertical integration of multi-omic single-cell datasets. We utilized this method to test our deep learning-based model against a matrix factorization algorithm.

Two more algorithms were included in the benchmark for extending the scope of comparison. TotalVI [13] was included since it is a deep generative framework based on VAE models. The algorithm allows us to assess potential performance improvements afforded by the novel features in SCPRO-VI. Lastly, we included Mowgli [11] which is a very recent algorithm that combines Nonnegative Matrix Factorization with Optimal Transport. Their study includes several comprehensive experiments demonstrating the efficiency of their algorithm against its competitors. Thus, we established a more comprehensive and contemproray comparison against state-of-the-art algorithms by including Mowgli.

Integration results were visualized using UMAP (Fig. 4A) and 1-NN cell matching, ASW, NMI, ARI, and graph connectivity metrics were employed for numerical comparisons (Fig. 4B). The UMAP projections provide insights into the solution approaches of algorithms and facilitate the annotation of numerical comparison results. SCPRO-VI and TotalVI were achieved to reduce intersections among different types of cell clusters by minimizing intra-cell distances. Seurat algorithm also increases the heterogeneity for some cell types like common lymphoid progenitors and lymphoid lineage-restricted progenitors. However, the algorithm separates cells of the same type into sub-clusters that are distantly placed from each another. This intra-cluster fragmentation may lead to poor or faulty downstream analyses are require comprehensive differentiation between different cell types.

**Fig. 4.**
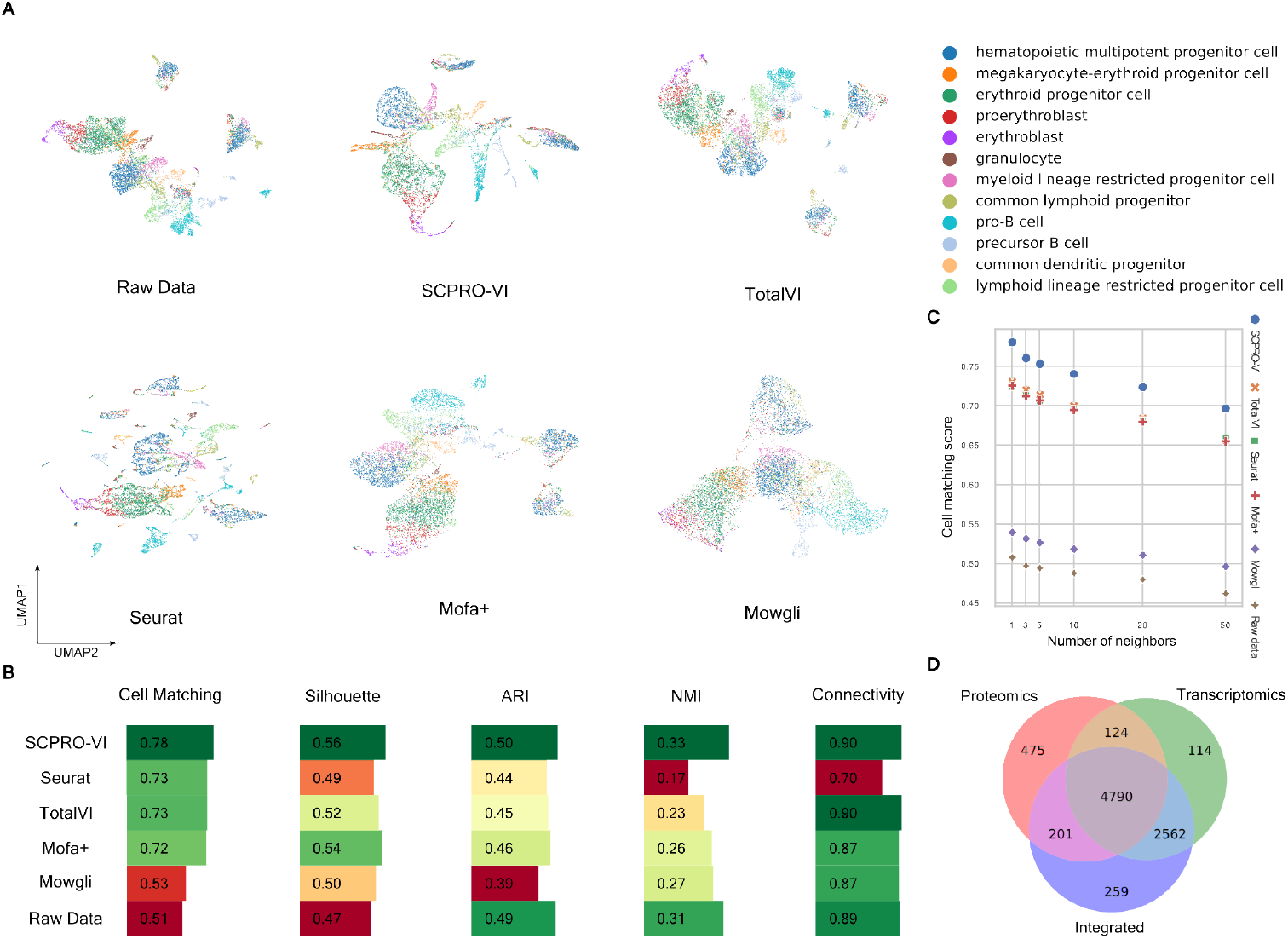
Benchmark results of algorithms on MNC dataset. **A** The UMAP projections of algorithm integrations colored by cell type. **B** Performance results of algorithms in numeric comparison metrics. **C** Purity analysis of algorithms in growing neighborhood sizes. **D** The Venn diagram of correct 1-nn cell matchings by transcriptomics, proteomics, and SCPRO-VI integrations.

Matrix factorization-based methods Mofa+ and Mowgli had similar outcomes with poor separation between different types of cell clusters. However, Mofa+ performed better than Mowgli in gathering the same type of cells together whereas Mowgli heavily mixed the cell clusters. On the other hand, none of the integration algorithms could combine the same type of cell clusters from different samples (see Supplementary Fig. 4A). However, only SCPRO-VI and Mowgli maintained to keep the precursor B cell clusters from different datasets together, which are weakly connected in raw data.

SCPRO-VI obtained the highest performance results in all the metrics against its competitors (Fig. 4B). Our algorithm improved the 1-NN cell matching score of raw data by 53%. The closest competitors TotalVI, Seurat, and Mofa+ had similar cell matching scores that are about 7% worse than our 78% matching score. Mowgli showed minimal improvement over raw data due to heavy intersection between cell clusters, as seen in its UMAP projection.

We also tested the quality of cell matching in growing neighborhoods (Fig. 4C). Results showed that all the algorithms experienced similar performance declines as the number of neighbors increased. However, it’s important to mention that when the 10 nearest neighbors were considered, SCPRO-VI had a similar matching performance with other algorithms’ best performances obtained for 1-NN matching. The pure local neighborhood led to higher performance results in other metrics since grouping similar cell types is essential for robust integration.

The silhouette score indicates the quality of inter-cell type heterogeneity, which is typically low in benchmark dataset due to the high intersection between different cell clusters. SCPRO-VI had the best performance by having 4% better results than Mofa+, the second-best algorithm with a 54% silhouette score. TotalVI had an average improvement by having a 52% silhouette score. Mowgli and Seurat had similar low results 50% and 49%, respectively. However, Seurat split the same cells into multiple sub-clusters which are distant as mentioned before. These distant cell clusters decrease the silhouette score of algorithms because the average silhouette score metric is calculated by considering one cluster center for each cell type (see Methods). Lastly, Mowgli had the lowest performance result due to the mixed cell clusters.

Our algorithm employed the best results in ARI and NMI metrics which demonstrate the alignment of cell clustering in SCPRO-VI embeddings with the ground truth cell clustering. SCPRO-VI is the only algorithm achieved to improve the performance of raw data by 2% whereas all other algorithms had worse alignment than raw data. The competitors except Mowgli exhibited similar performances. Mowgli distinguished with the lowest ARI score since the algorithm failed to exclude different types of cells. Moreover, Seurat had the lowest NMI score due to the intra-cell type fragmentation. Graph connectivity scores are very close for all benchmark algorithms except Seurat due to the same reason for having low NMI scores. SCPRO-VI and TotalVI generated 1% better embeddings than raw data by having a 90% graph connectivity score which means the same type of cells were gathered closer to shape-connected clusters than raw measurements. Mofa+ and Mowgli had slightly worse connectivity scores with 87% since they succeeded in embedding the same type of cells together.

Benchmark results demonstrate the efficiency of the SCPRO-VI algorithm for multi-omic data integration. Our algorithm surpassed state-of-art comparison algorithms at five performance metrics to measure the heterogeneity increment between different types of cell clusters while decreasing intra-cluster distances of cells in the same type. Thus, we showed that the SCPRO-VI algorithm is the best algorithm among the benchmark algorithms for meeting the two fundamental aims of multi-omic data integration.

SCPRO-VI integration was compared also with single modalities to calculate the contribution of integration to the cell matchings (Fig. 4D). 1-NN neighbors were calculated using cosine distances between cells by measurements of each modality. Of note, we used PCA embeddings instead of raw measurements while calculating distances by transcriptomics (see Methods). PCA embeddings increased the matching performance dramatically from 54% to 76%.

Results showed that our algorithm leverages both modalities to outperform individual modalities. Moreover, we observed that transcriptomics has more inter-cell type heterogeneity than proteomics for each cell type in the benchmark dataset since there are measurements from only 176 proteins against 11361 genes (see Supplementary Fig. 4B). The results also showed that 5% of the cells (475 cells) were matched truly by proteomics but mismatched by both transcriptomics and integrated distances. For example, 143 EPC cells and 135 hematopoietic multipotent progenitor cells were truly matched by their protein abundances despite the number of cells only matched truly with transcriptomics are 24 and 29, respectively.

These results indicated that transcriptomics dominated proteomics in SCPRO- VI integrations when they are conflicted. However, SCPRO-VI effectively managed the disagreements and achieved to increase the aggregation of modalities from 49% (4790 + 124 cells) to 75.5% (4790 + 201 + 2562). Moreover, SCPRO-VI successfully matched 2.6% cells which are mismatched by both proteomics and transcriptomics measurements.

### 2.6 Analysis of integrated cell embeddings assessing increased heterogeneity and revealing custom cell sub-clusters

We assessed our novel algorithm’s efficiency and effectiveness by further downstream analyses on benchmark dataset to see its capabilities. First, we performed a density-based clustering with DBSCAN [29] algorithm on integrated embeddings of SCPRO-VI for revealing distinguished cell sub-clusters (see Methods). Among 18 cell clusters with more than 50 cells, we selected one granulocyte, and one common lymphoid progenitor (CLP) cell clusters which are pure sub-clusters distinguished from the main clusters of their cell types.

The two cell sub-clusters are browsed in the clusterings of other algorithms’ integrated embeddings. In this analysis, we aimed to compare the capability of our algorithm against other integration methods in the discovery of sub-clusters that aren’t detectable in raw data. The granulocyte cell cluster is easier to detect than CLP cells, as they are nearly separable even in raw data. On the other hand, the cells in the CLP sub-cluster are contained at the center of the main CLP cluster without any separation. We selected this sub-cluster because of its more challenging nature than the granulocyte cells sub-cluster.

We calculated the optimal clusterings of algorithms for both cell sub-clusters individually (Fig. 5A). We systematically calculated several clusterings for each algorithm and ranked them by an F1 matching score (see Methods). Each cluster in a clustering is matched with the CLP cell sub-cluster, and the F1 score is calculated by the co-occurrences of cells in clusters. The clustering containing the cell cluster with the highest F1 score was determined as optimal. The same procedure was applied for the granulocyte sub-cluster, thus two optimal clusterings were obtained for each algorithm (Fig. 5A).

**Fig. 5.**
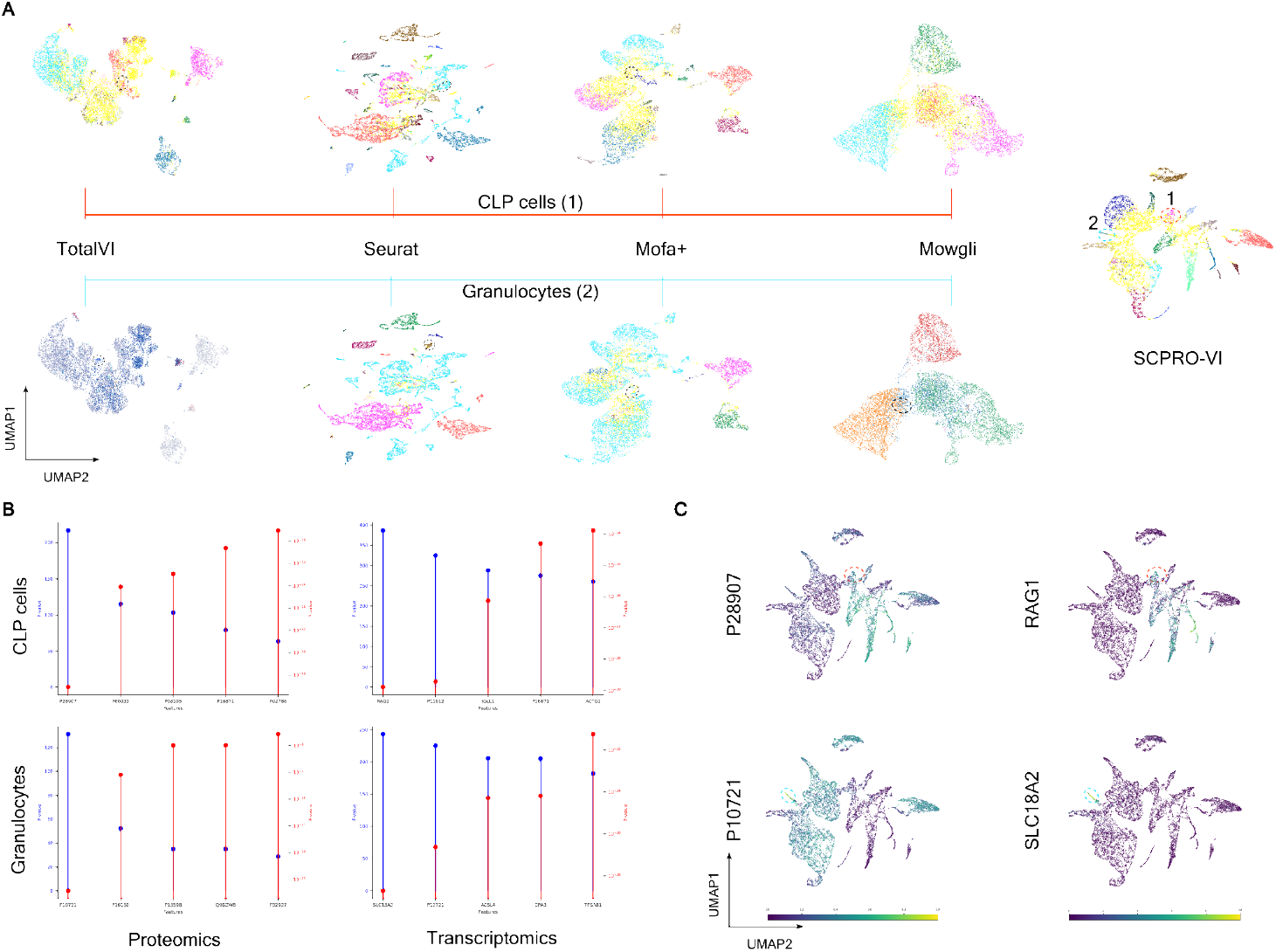
The results of novel cell sub-cluster analysis in integrated MNC data. **A** The UMAP projections of integrated embeddings of algorithms colored by own optimal clusterings of each algorithm. The optimal clusterings are calculated and projected individually for both CLP (top) and granulocyte (bottom) sub-clusters. The cells in the target sub-cluster are circled. **B** The Stem diagram of *P* and *F* values of important proteins (left) and important genes (right) for both CLP (top) and mono- cyte (bottom) cell sub-clusters. **C** The UMAPs of integrated embeddings of SCPRO-VI colored by the expression level of most distinctive protein (left) and gene (right) for both CLP (top) and mono- cyte (bottom) cell sub-clusters.

The results for the granulocyte cells show that the Seurat algorithm achieved to cluster of these cells in a nearly pure cell cluster by 0.82% F1 score. Of note, our algorithm F1 scores of both granulocyte and CLP cell clusters are 1 since they are clustered in separated cell clusters containing no other cell. Seurat’s success is expected since it tends to split the cells into pure small clusters. TotalVI has also achieved a good performance by having a 76% F1 score. This VAE-based method reveals the power of deep learning in capturing small variations in cell clusters while preserving their inter-cell heterogeneity. Mowgli and Mofa+ couldn’t distinguish the granulocyte sub-cluster from the rest since both generated intersected cell clusters (F1-score 0.02% for both algorithms).

In order to examine examine the biological significance of the granulocyte cell subset, we conducted distinctive gene and protein analyzes (Fig. 5B, see Methods).

The P10721 gene and protein, which are frequently expressed in granulocyte cells, appeared as intra-cell type discriminators, suggesting that this cell subset differentiated into basophil, natural killer (NK), or eosinophil cells [30]. However, upregulation of the SLC18A2 gene from other granulocyte cells (Fig. 5C) increases the possibility of being basophil cells [31]. In addition, distinctive proteins P16150 (CD43), P13598 (CD102), and Q9BZW8 (CD244) support the possibility of being basophil cells regarding their proteomic measurements [30, 32]. In conclusion, we claimed that the detected sub-cluster is actually specialized for a specific task and our algorithm is capable of revealing such sub-clusters.

We repeated the same procedures for the CLP cell sub-cluster to evaluate the comparison algorithms with a sub-cluster that is more difficult to detect. Mowgli and Mofa+ algorithms failed to detect as expected since they couldn’t detect the granulocyte sub-cluster. TotalVI couldn’t maintain its success while revealing CLP sub-cluster by having 0.16% F1 score. Seurat algorithm performed as the second best algorithm with 70% success. However, we observed that all the CLP cells were clustered into one big cluster which means that Seurat is capable of distinguishing CLP cells but couldn’t distinguish the targeted sub-cluster from the rest. All these results indicate that SCPRO-VI is capable of finding rare cell sub-clusters that emerged in integrated data that couldn’t be revealed by other multi-omic data integration algorithms.

Distinctive genes and proteins in CLP sub-cluster were queried as done in granulocyte cells (Fig. 5B). Identified distinctive genes RAG1, P11912, P16871, ACTG1, IGLL1 cause differentiation and formation of sub-clusters in common lymphoid progenitor cells [33–35]. Similarly, identified distinctive proteins P28907, P60033, P16871, P08195, and P26010 have been reported in the literature for the contribution to cell differentiation in CLPs [36, 37]. The most distinctive gene RAG1 and the most distinctive protein P12907 upregulated in these cells unlike the other CLP cells (Fig. 5C). However, the upregulation of both RAG1 and P12907 isn’t unique for the cells in the identified sub-cluster.

We employed highly variable gene/protein (HVG / HVP) analysis using the Seurat v3 [38] algorithm for a better annotation of the CLP sub-cluster. The most variable five genes ZNF300, ITGB7, LINC01954, DIABLO, EHD3, and five proteins P49961, IgG2b, ICOSLG, SELL, Q9BQ51 were identified in the HVP analysis. It is reported that the IgG2b shows significant expression changes during the differentiation of CLP cells [39]. Moreover, co-existing highly variable changes of IgG2b, ICOSLG, PDCD1LG2, and SELL proteins together suggest the differentiation of CLP cells into B cells [39, 40]. Among these proteins, only IgG2b was got into the 10 most HVPs of all CLP cells. This indicates that this specialized cell cluster has its own HVGs. All these results suggest the effectiveness of our algorithm in multi-omic data integration by generating strong multi-omic embeddings to unlock rare cell populations.

## 3 Methods

### 3.1 Dataset preparation

In this study, we used two multi-omic CITE-seq datasets [3, 20]. Both datasets were obtained from the Human Cell Atlas (HCA) database [41] in annData format. The datasets were converted into MuData format, an extended version of annData for handling multi-omic datasets [42]. The features were split into transcriptomics and proteomics by referencing the metafiles from their HCA repositories, as the original annData files lack explicit feature type information.

The HIV [3] dataset (*Gene Expression Omnibus accession ID: GSE164378*) comprises 161764 cells from 8 HIV patients, sampled at three intervals: before vaccination, 3 days after vaccination, and 7 days after vaccination. There are measurements for 188 proteins and 20380 genes in the dataset. We separated the cells into 24 datasets by patient and sampling time. We also tested our algorithm performance in a multi-sample dataset sampled randomly from all 24 datasets. We selected 5k cells in total which aligns with the cell counts in the individual datasets. The number of cells from each patient is proportional to the original dataset (see Supplementary Fig. 5A).

The MNC dataset contains measurements of 197 proteins and 31770 genes obtained from 28k cells. TotalVI and Mowgli couldn’t be tested on the full dataset in our experimental setup due to their high computational power requirements. Therefore, we randomly sampled 10k cells from the dataset without regard for any specific dataset characteristics. The new dataset contained an equally proportional number of cells for each cell type as in the original dataset (see Supplementary Fig. 5B).

### 3.2 Data preprocessing

All datasets were processed without any normalization since the HIV dataset was already log-transformed and the MNC dataset contains normalized data both with positive expression values for all features. Principal component analysis (PCA) was applied to the transcriptomic data to increase inter-cell heterogeneity while suppressing the noisy measurements causing conflict. The number of components is set to 100 which is coherent with proteomics dimensions and explains data more than 90% for both datasets.

The genes in transcriptomic data are filtered by the number of cells they are measured. We set a default threshold 5% in our experiments which can be adjusted with a trade-off between trusty or comprehensive feature set. The number of genes decreased to 8417 for the subset of the HIV dataset. The same filtering procedure decreased the included gene number to 11361 in the MNC dataset. On the other hand, we used all protein measurements since the number of sequenced proteins is limited.

### 3.3 SCPRO-VI

Our integration method consists of three main steps: calculation of distances between cells by each modality, pre-training of modality embedding models, and integration of modality embeddings (see Fig. 1). First the algorithm generates cell-to-cell distance matrices individually for each modality. The distance matrices are filtered then to calculate the adjacency matrix used in VGAE models. The VGAE models embed each modality into a discrete data space of matching dimensions. Finally, the modality embeddings are integrated by the joint homogeneous fusion approach [17] with an encoder model that is connected to the output of VGAE models and generates the integrated cell embeddings as output. All steps of the SCPRO-VI algorithm are detailed in the following subsections.

#### 3.3.1 Distance matrix calculation

We proposed a novel metric for measuring cell-to-cell distance. Let *P*_1_ be the Hadamard product of feature matrix *A*_1_ of *cell*_1_ with its transpose *A*_1_*^T^* and *P*_2_ is the same product for *cell*_2_. The distance between *cell*_1_ and *cell*_2_ is calculated as |(*P*_1_ −*P*_2_)∗*W* |*/*(|*A*_1_|∗|*A*_2_|) where *W* is the matrix of importance values between all feature pairs. The resulting sum is normalized by the *L*_1_ norm multiplication of cells, |*A*_1_| and |*A*_2_|, for avoiding expression level differences and focusing on the co-expression profiles of the cells. The metric is formulated as below:

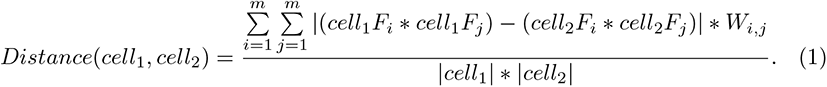

Where, *F_i_*and *F_j_*are the i-th and j-th features in the related cell. *W_i,j_* is a weighting factor indicating the relevance score between feature *i* and *j*. The *L*_1_ norms of cell vectors are represented with |*cell*_1_| and |*cell*_2_|. The distance metric may also be considered as the isomorphism score between two weighted complete graphs of cells where the nodes are features and edges are the product of features weighted by relevance scores.

Protein-protein interaction values are used as relevance scores in distance matrix calculations by proteomic data. The weights are obtained from the STRING [19] database proposed as association scores (v12.0). The scores are between 0 and 1 and are used as they are in the experiments. Moreover, we set the relevance score to 1 for the self-pairs of features.

Cosine distance was applied in place of the novel distance metric for calculating cell distances by transcriptomics for three reasons. First, we use PCA embeddings instead of the raw data which disabled the use of our metric since there isn’t any biological relevance between components. However, we still tested the algorithm by raw measurements to see its feasibility in other omic types. We used pathways for calculating the relevance score. The pathways are obtained from the Reactome [43] database at all levels of the pathway hierarchy. The relevance score between two genes was calculated as the number of co-occurrences of genes in the same pathway. The occurrences are min-max normalized before using them as the weights in distance calculation.

The results showed that the relevance scores increased mismatches from minority cell types with fewer members to the majority cell types such as monocytes. We observed that the longer pathways led to increased relevance scores between commonly expressed genes. As a result, the distinctive unique co-expressions causing differentiation of rare cell clusters were suppressed by low relevance scores. As a result, we discontinued the use of pathways for biological relevance scores between genes. We also didn’t set up any further experiments for calculating cell distances by transcriptomics using the novel distance metric.

The last but not least reason is the complexity of our distance metric, though its parallelizable algorithm. The Hadamard multiplications are done fully parallel since they are distinct vectorial multiplications for each cell. Moreover, the subtractions are done in parallel and can be batched concerning memory consumption. However, the complexity of the distance metric is *O*(*n*∗*n*∗*m*∗*m*) where *n* is the number of cells and *m* is the number of features. The complexity of the algorithm causes high computational needs since the complexity quadratically increases both by the number of cells and by the number of features. Therefore, it makes it less desirable for transcriptomic data with more than 10k features.

We normalized the distance matrices to avoid domination of modalities in integration raised by scale imbalance between them. For each modality, each cell normalized individually by dividing the mean distance of the nearest 100 neighbors. Thus, we get the distance ratios instead of modality-specific distance values for the nearest 100 neighbors of cells.

The number of nearest neighbors *N* considered for normalization declares the considered intra-cell neighborhood size for the cells. Increasing the neighborhood size may raise again the burden caused by scale differences. However, narrow neighborhoods can also lead to this issue since the normalization affects only a few intra-cell type distances and the rest stay unaffected.

Hadamard multiplication is applied to these normalized distance matrices to yield the integrated distance matrix *D^I^* . These integrated distances are then used in the training of end-to-end multi-omic data integration model.

#### 3.3.2 Embedding modalities

Transcriptomics and proteomics were embedded independently using distinct VGAE models for better convergence while training the end-to-end multi-modal integration model. Pre-training of the embedding models enhances the inter-cell type heterogeneity within each modality (see Supplementary Fig. 6A). Each model was trained on its respective modality distance matrices. These pre-trained models transfer modality-specific cell neighborhoods into the integration model. This approach improved convergence during end-to-end training, allowing the model to focus primarily on integration rather than on learning cell relations within individual modalities.

Each VGAE model receives raw measurements for each modality, paired with the modality-specific graph, and outputs a vector representing the cell embedding (see Supplementary Fig. 6B). The modality-specific graph is constructed through filtering of the modality’s distance matrix. We propose the following filtering function to generate the adjacency matrix:

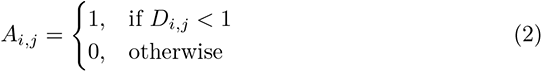

Where, *A* represents the adjacency matrix filtered from the modality cell distance matrix *D*. *A_i,j_* and *D_i,j_* denote the adjacency and distance relationships between cells *i* and *j*, respectively. This equation enables identification of each cell’s *k* nearest neighbors (where *k* ranges from 0 to *N*) as entries in the adjacency graph. We applied a constant threshold value of 1 for filtering, given that the distance matrices were normalized by the mean distance of each cell’s *N* nearest neighbors. For each cell, *k* neighbors had distances below this mean, while all other distances exceeded 1. This consistent threshold value across datasets offers an adaptive neighborhood size per cell, avoiding the need for hyperparameter optimization. Adaptive neighborhoods help mitigate mismatched neighbors often seen in fixed k-nn approaches, allowing precise handling of cells within non-convex regions without overly restricting neighborhoods for cells within central cell clusters (see Supplementary Fig. 6C).

The variational graph auto-encoder model comprises an encoder alone, as it is optimized to learn neighborhood structure rather than to reconstruct inputs. The following loss function computes the mean element-wise Manhattan distance between the actual modality adjacency matrix and the predicted adjacency matrix, separately accounting for positive and negative relations:

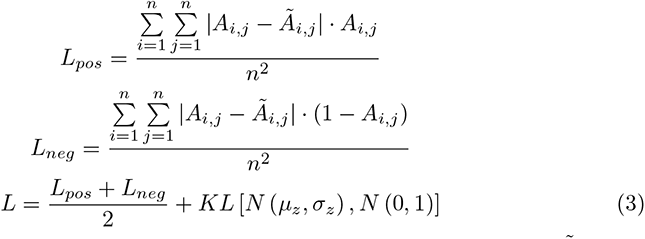

Where, *A* represents the adjacency matrix for a given modality and *Ã* is the adjacency matrix predicted by the model. *KL* donates the Kullback-Leibler (KL) divergence [44], which measures the discrepancy between the distribution of the sampling function in the latent space and a standard Normal distribution.The KL divergence regularizes the latent space, encouraging it to conform to a structured distribution. *Ã* is calculated as follows:

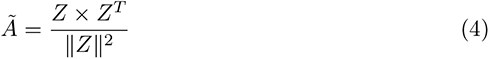

Where, *Z* is the matrix of cell latent space embeddings. Values in *Ã* range between 0 and 1, with higher values assigned to cells with more similar embeddings. This enables the training of VGAE models by reducing similarity among irrelevant cells while enhancing similarity among neighboring cells. The contrastive learning approach, which individually treats positive and negative samples in the loss calculation, mitigates the dominance of negative samples typically due to the sparsity of similarity graphs.

The linear model employed for comparisons has the same model architecture with VGAE model, but it replaces graph convolution layers with linear layers (see Supplementary Fig. 6C). Thus, the embedding model turns into an encoder of a vanilla VAE model. The loss function in Equation 3 is applied without the Kullback-Leibler divergence term.

UMAP embeddings of integrated distance matrices were calculated for comparison. The UMAP embeddings were generated via the *tl.umap* method within the Scanpy package [45]. We set the integrated distance matrix as the *connectivities* parameter.

The *distances* parameter was assigned a filtered integrated distance matrix, with only the 30 nearest neighbors of each cell retained and all other distances set to 0. The considered number of nearest neighbors is set to 30 as recommended in the *RunUMAP* function within the Seurat package. All remaining parameters were kept at their default values.

#### 3.3.3 Multi-modal integration

We developed a multi-omic integration pipeline using a multi-view variational graph auto-encoder (VGAE) model followed by a 2-layer encoder for modality fusion (see Supplementary Fig. 7A). Modalities are independently embedded using modality- respective pre-trained VGAE model. Each VGAE model receives raw modality measurements along with the modality-specific similarity graph, generating latent embeddings for each cell. The cell embeddings are concatenated and provided as input to the encoder. The modalities are fused into an integrated latent space as model output.

The end-to-end model is trained using a loss function similar to that used in training the VGAE models (see Eq. 3). However, the KL divergence for both modality distributions was incorporated to avoid distortion in modality embeddings caused by overemphasis on a single modality. We obtained the ground truth adjacency matrix by filtering the integrated distance matrix *D^I^*with the equation below:

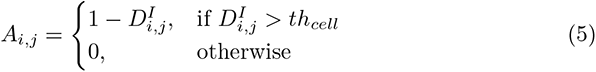

Here, *A_i,j_* and *D^I^* represent the adjacency and distance relation between cell *i* and *j*, respectively. The threshold *th_cell_*filters out irrelevant relations between cells that are not in the considered neighborhood. This threshold is calculated as the mean distance to each cell’s 100 nearest neighbors. The number of cells in the designated neighborhood was set to 100, as specified in the distance matrix calculation section.

Filtered distances are converted to a weighted adjacency matrix by subtracting them from 1 as in the equation 5. The predicted adjacency matrix *Ã* is calculated by applying the equation 4 over integrated embeddings. Eventually, not only 0-1 relations between cells are considered as in the modality embedding models, but also the degree of relations are optimized with the loss function. Therefore, pruning about 99% of relations that are unimportant for integration enhances the convergence of the end-to-end model. Moreover, the convergence speed is also increased by filtering since the optimizer struggles with fewer values different than 0.

The contrastive learning approach is also employed in the loss function of the end-to-end model. The positive and negative samples are handled individually and mean element-wise Manhattan distances are calculated separately similar to the Equation 3. However, we calculated the unweighted adjacency matrix *A^U^* for masking, since *A* contains edge weights instead of 0-1 relations. *A^U^* is generated by applying Equation 2 to distance matrix *D^I^* . Thus, we are able to filter element-wise Manhattan distances only between positive samples by multiplying the calculated distances with *A^U^* . The negative loss is calculated by multiplying the distances with 1 − *A^U^* as in the Equation 3.

#### 3.3.4 Environment and model setup

The embedding and integration methods were implemented using the PyTorch ecosystem (version 2.4.0). In the VGAE model, we used GraphSAGE [46] layers as graph convolution layers. The number of neighbors was set to 20 for the first convolution and 10 for the second convolution for all the layers to be coherent with the original study. The first convolution layer dimension is 256, and the following *µ* and *σ* convolution layers are of the same size with latent embeddings in 128 dimensions. We set the dimension of latent space by considering the input size of each modality. The encoder consists of a layer with 512 dimensions followed by another layer in half size. The integrated embeddings are also in 128 dimensions.

We used Adam optimizer (*lr* = 0.001 as default) in the optimization of all models. The models were trained in 500 epochs in the experiments by default. However, we observed that the VGAE models converged mostly after 200 iterations but the loss value decreased steadily until 1000 epochs (see Supplementary Fig. 7B). Moreover, the end-to-end model converges even faster in 50 epochs but the improvement stands still until 2000 epochs. We didn’t employ any further hyperparameter optimization in our experiments.

### 3.4 Benchmark algorithms

#### 3.4.1 Seurat v4

We used Seurat v4 [3] in our comparisons. The Seurat R package was used for implementation. We used weighted-nearest neighbor distances while calculating *k*-nn cell matching scores. All other metrics were calculated using the UMAP embeddings obtained using the *RunUMAP* method with default parameters.

#### 3.4.2 TotalVI

We used the TotalVI [13] deep learning algorithm in our benchmark on the MNC dataset. The algorithm couldn’t be tested in experiments done with the HIV dataset because the algorithm operates only with unnormalized protein counts which is not available for this dataset. We used the implementation of the algorithm provided in the scVI [22] package with default parameters.

#### 3.4.3 Mofa+

We used the Mofa+[9] algorithm in our benchmarks since it is one of the most common comparison algorithms for multi-omic data integration [47]. We tested the algorithm using the implementation in the muon package which is a wrapper of the mofapy2 package. All the parameters are used as defaults in muon implementation.

#### 3.4.4 Mowgli

We compared our results with Mowgli [11], which is another matrix factorization algorithm optimized by optimal transport. The algorithm requires highly variable features for integration. We calculated the highly variable genes using the Seurat v3 [38] package. We set the number of HVGs to 2000 as in the Seurat v3 paper. Moreover, the authors of Mowgli used 2500 HVGs for about 35k genes and 1500 HVGs for about 8k genes. Thus, we decided to use 2000 HVGs for about 12k genes in our filtered benchmark dataset. We set all proteins as highly variable as in the original study.

### 3.5 Metrics

The average silhouette width, adjusted rand index, normalized mutual information, and graph connectivity metrics are calculated using the Scib package [48] with default parameters. K-nearest neighbor cell matching score and F1 score were implemented in our package.

#### 3.5.1 Cell matching

The cell matching score, or k-nn matching score, measures the purity of local neighborhoods of cells by the ground truth cell types. The number of truly matched neighbors of each cell is calculated and averaged by the following equation:

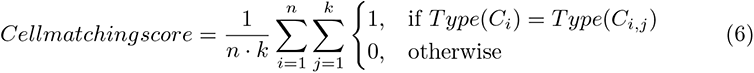

Where, *C_i_*is the i-th cell, *C_i,j_* is the j-th nearest neighbor cell of the i-th cell, and *Type*() is the ground truth labeling function. Thus, if all the cells truly match with k-nearest cells, then the metrics become 1 and it takes 0 when there aren’t any neighbors in the same type for none of the cells.

#### 3.5.2 Average silhouette width (ASW)

The average silhouette width metric reflects the degree of both intra-cluster similarity and inter-cluster heterogeneity at the same time. The metric is calculated by measuring the silhouette coefficient of each cell *c* by using the (*b* − *a*)*/max*(*a, b*) equation. Here, *a* is the mean distance of cells in the same type with *c* to the cluster center and *b* is the mean distance to the center of all other cell type clusters. The values of ASW are between 1 and -1 where 1 is the best score.

The Scib package follows the same implementation as the Scikit-learn package [49]. The values are normalized in Scib implementation by (*ASW* + 1)*/*2 equation to scale them between 0 and 1. The ASW values around 0.5 indicate gathering clusters by cell types but still, there are heavy overlapping among the clusters. The ASW values close to 0 means that there is no significant clustering among the cells in the same type and the whole data is a mess.

#### 3.5.3 Normalized mutual information (NMI)

Normalized mutual information is a measurement of the alignment between a ground truth clustering schema and computational clustering schema of the data. The value of NMI metrics is between 0 and 1 where 1 indicates a perfect matching between ground truth cell clusters and the clustering in integrated data. The uncorrelated clusterings resulted in have low NMI score. Therefore, higher NMI scores demonstrate the intra-cell type gatherings in integrated data.

We used the *nmi*() function of the Scib package with default parameters in our experiments. This method gets the ground truth cluster labels and generates a clustering using the Leiden algorithm [50]. We used ground truth cell types as the cluster labels, and the Leiden algorithm calculated an optimized clustering by the given cell type labels. Scib uses the *normalized mutual info score* function of the Scikit-learn package with an arithmetic averaging method to calculate the NMI score.

#### 3.5.4 Adjusted rand index (ARI)

The adjusted rand index metric calculates the alignment of two clusterings as NMI by counting the cell pairs that co-existed in aligned cell clusters. However, ARI also considers the common pairs between the cells in different clusters. Thus, the ARI metric reflects the differentiation of the different types of cell clusters which is not the case in the NMI metric.

The ARI metric is calculated by the *ari* function in the Scib *Metrics* module. The method uses the *adjusted rand score* function from Scikit-learn which takes ground truth labels and predicted labels as inputs and calculates the ARI score. We used the cell types as ground truth labels and predicted clustering labels of the Leiden algorithm as explained in the NMI metric. The metric gets values close to 0 when there is no matching between clusterings. In our case, it means the integration resulted in random embeddings with no similarity between the same type of cells. The performance results close to 1 indicate high intra-cell type clustering along with the inter-cell type differentiation.

#### 3.5.5 Graph connectivity

Graph connectivity score measures how the nodes of the same type are connected together. The following equation is used in Scib for calculating the connectivity metric:

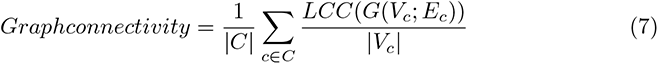

Where, *C* is the set of unique cell types, *c* is each unique cell type and *V_c_* is the set of nodes in *c* cell type. *LCC* is a function that gets a graph as input and returns the number of nodes in the largest connected component in the input graph. *G*(*V_c_*; *E_c_*) is a graph where *V_c_* is the set of nodes as the cells in type *c* and *E_c_* is the set of edges between these nodes. Thus, *LCC*(*G*(*V_c_*; *E_c_*)) is the number of nodes in the largest connected components of cells in *c* cell type. Consequently, the equation calculates the ratio of cells in connected components to the cells in each cell type and averages the maximum ratios obtained for each cell type.

The graph connectivity metric is useful for evaluating how the cells in the same type are similar in integrated data. Moreover, this metric is sensitive to intra-cell type fragmentation which leads to lower connectivity scores. On the other hand, the metric performs individual calculations for each cell type. Thus, such an integration without any inter-cell heterogeneity but with strong intra-cell similarity leads to high-performance results. Therefore, the metric should be evaluated with other performance metrics while assessing the overall performance of integration.

#### 3.5.6 F1 score

The F1 score was used to calculate the alignment of a query cluster with clustering in our experiments. The F1 score is calculated by the following equation:

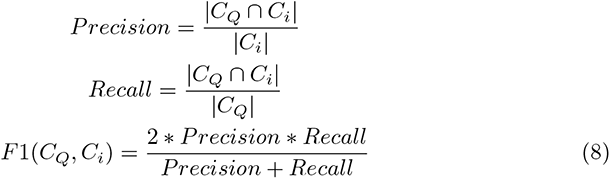

Where, *C_Q_* is the set of cells in the query cluster and *C_i_* is the set of cells in the i-th cluster of a clustering. We calculated the rank of a clustering as *MAX*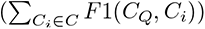. The equation ranks the clusterings by the maximum F1 score between the query cell cluster and their cell clusters. Thus, we were able to identify the best clusterings of algorithms that have the potential to reveal analyzed sub-clusters.

### 3.6 Data analysis

#### 3.6.1 Distinctive feature analysis

Distinctive proteins/genes are determined using the *f classif* method from the Scikit-learn package. This method takes features and target labels as input and calculates univariate feature scores by applying an ANOVA F-test to each feature for the classification in labels. We set the labels of cells in the same type to 1 and the rest of the cells to 0 while calculating the distinctive features of a cell type. On the other hand, the distinctive features of novel cell sub-clusters are calculated using only the cells in the same type as the query sub-cluster. We set the labels of cells in the sub-cluster to 1 and the rest of the cells to 0. The features are sorted by their F-test scores and the top 5 features were identified as distinctive features.

#### 3.6.2 Cell sub-cluster identification

Density-based clustering algorithm DBSCAN [29] was used in sub-cluster analysis since the single-cell datasets usually exhibit non-convex cell clusters. We used the Scikit-learn implementation of the DBSCAN algorithm. The number of minimum samples was set to 5 as default which is coherent with the minimum number of cells in each cell type. In the tests over the MNC dataset, we employed a systematic search for selecting the best eps value for each algorithm. First, we sorted the distances of cells to their 5th nearest neighbors since the minimum number of considered neighbors was set to 5. The minimum and maximum distances were identified, and the 20 equally spaced distances were enumerated and each was used as the eps value for an individual clustering. The number of eps values is set to 20 for employing 5% exchange rate resolution. The number of eps values can be increased by dividing more chunks for the more complex datasets.

The systematic clustering was done first for the SCPRO-VI algorithm to identify novel cell sub-clusters. We filtered the cell clusters with more than 50 cells and consisted of cells all of the same type. CLP and granulocyte cell sub-clusters are identified in the same clustering with *eps* = 0.005 valued. Then, the process was applied for each algorithm and 20 different clusterings were obtained for each of them. The clusterings were ranked for both CLP and granulocyte cell sub-clusters by individual F1 scores. The clustering or clustering pair with the highest rank for each sub-cluster was determined as the optimal clustering for each algorithm.

We followed a similar procedure in the identification of monocyte cells in the targeted sub-cluster revealed on patient 1 day 7 samples in the HIV dataset. The number of minimum samples was set to 5 and a systematic search was applied by the 20 different eps values. The resulting clusterings were evaluated by counting the number of cells contained by the revealed monocyte sub-clusters in the targeted region of the monocyte cluster. We identified the largest monocyte sub-cluster in our embeddings with *eps* = 0.03 valued.

#### 3.6.3 Gene ontology (GO) analysis

Gene ontology analysis was done individually for the most important and least important proteins in each dataset. The most and least important proteins in a dataset were identified by the protein-protein interaction scores. The PPI pairs were filtered first for the proteins in the dataset and then sorted by their interaction scores. Proteins appeared in the top 50 interactions identified as most important proteins, and the proteins appeared in the last 50 interactions identified as least important proteins.

We used g:Profiler [51] for annotating the most and least important proteins in two separate rounds. The default parameter setup was used in the analysis. The results were listed in detail for each annotation term: molecular function, cellular component, and biological process (see Supplementary Fig. 8).

## 4 Discussion

Integrated analyses of omic layers in single cells enhance cellular granularity, enabling the identification of specialized cell subgroups. However, encoding relationships from two distinct data spaces into a joint space remains challenging due to conflicts arising from imbalanced omic-specific relationships. Furthermore, effective integration must also be explainable and interpretable to provide deeper insights into regulatory mechanisms across omic layers.

In this study, we introduce Single-Cell PROteomics Vertical Integration (SCPRO- VI), a novel tool for integrating multi-omic single-cell data, specifically focusing on paired proteomic and transcriptomic datasets. Notably, SCPRO-VI is capable of integrating any paired modalities with numerical features, even though this research primarily concentrates on proteomic and transcriptomic integration. The algorithm computes integrated distances among cells by multiplying balanced omic-specific distances, calculated using a novel biologically aware distance metric. Subsequently, SCPRO-VI employs a multi-view variational graph auto-encoder to embed these modalities into a joint data space, accurately reflecting the relationships captured in the integrated distances.

We rigorously tested SCPRO-VI using a multi-sample CITE-seq dataset [3], evaluating its robustness, efficiency, and effectiveness. The results demonstrated superior performance compared to competing methods across all 24 sample datasets, based on computational metrics for cell type heterogeneity and subgroup detection. Importantly, our analyses revealed that SCPRO-VI amplified intra-cluster heterogeneity, uncovering biologically meaningful sub-clusters that are consistent with findings in the literature.

We further benchmarked SCPRO-VI against state-of-the-art algorithms using a more challenging CITE-seq dataset [20] characterized by lower inter-cell type heterogeneity. The results clearly show that SCPRO-VI outperforms existing methods in both computational metrics and its ability to reveal specialized cell sub-clusters. Additionally, SCPRO-VI preserved the purity of cell matching within expanding neighborhoods more effectively than its competitors, leveraging both modality-specific relationships.

It is important to note that the performance of state-of-the-art algorithms can vary significantly across different datasets, as observed in our experiments. These comparisons were conducted using the default parameter settings recommended by the respective algorithm authors, which may limit generalizability. Furthermore, the reliance on cell type annotations, often derived computationally by the dataset creators, introduces potential inaccuracies due to mislabeling. While all algorithms were evaluated under the same annotation framework, mislabeled cells may have adversely affected performance metrics.

One notable limitation of SCPRO-VI is the computational demand of its novel distance metric calculation, which has quadratic complexity. This poses challenges for datasets with millions of cells or thousands of features per modality. However, this limitation can be mitigated through parallelization, such as batching cells and features for efficient vectorized operations on GPUs. Another challenge is the lack of comprehensive ranking information for feature relationships. For instance, protein-protein interaction data are widely available for proteomics, but equivalent gene-gene interaction databases remain underdeveloped, despite progress in individual studies [52].

Several areas for improvement could further enhance SCPRO-VI. Currently, integrated distances are computed by simply multiplying balanced distances for simplicity. Developing an advanced algorithm that dynamically weights omic-specific distances based on their reliability could improve performance. Additionally, the introduction of an adaptive neighborhood size, determined through a preprocessing step, could optimize distance calculations and similarity graph generation, potentially boosting algorithm performance.

Beyond these extensions, one of the most exciting future directions involves addressing missing modalities. Compared to scRNA-seq datasets, the number and scope of multi-omic studies remain limited. Enhancing the embedding model with a generative module trained on multi-omic data could enable integration even when proteomic data are absent. Such an extension would allow SCPRO-VI to project multi-omic heterogeneity onto comprehensive single-cell transcriptomic datasets, broadening its applicability and impact.

## Data availability

The data analyzed in this study were obtained from the Human Cell Atlas (HCA) database [41]. The HIV dataset is accessible at https://explore.data.humancellatlas.org/projects/3ce9ae94-c469-419a-9637-5d138a4e642f, and corre- sponding meta-data available at Gene Expression Omnibus (GEO) database [53] under accession number GSE164378. Mononuclear cell data can be accessed at https://explore.data.humancellatlas.org/projects/04ad400c-58cb-40a5-bc2b-2279e13a910b, with metadata deposited in GEO under accession number GSE166895. Protein - protein interaction scores were downloaded from the STRING database [19] at https://stringdb-downloads.org/download/protein.links.v12.0/9606.protein.links. v12.0.txt.gz. Pathway data were obtained from the Reactome database [43] at https://reactome.org/download/current/UniProt2Reactome All Levels.txt.

## Code availability

The Python code of reference implementation and code to reproduce the results in this manuscript is available at https://github.com/single-cell-proteomic/SCPRO-VI.

## Notes

### Competing Interest Statement

The authors have declared no competing interest.

## References

[1] Cao, Z.-J. & Gao, G. Multi-omics single-cell data integration and regulatory inference with graph-linked embedding. Nature Biotechnology 40, 1458–1466 (2022). Publisher: Nature Publishing Group.

[2] Baysoy, A., Bai, Z., Satija, R. & Fan, R. The technological landscape and applications of single-cell multi-omics. Nature Reviews Molecular Cell Biology 24, 695–713 (2023). Number: 10 Publisher: Nature Publishing Group.

[3] Hao, Y. et al. Integrated analysis of multimodal single-cell data. Cell 184, 3573–3587.e29 (2021).

[4] Rautenstrauch, P., Vlot, A. H. C., Saran, S. & Ohler, U. Intricacies of single-cell multi-omics data integration. Trends in Genetics 38, 128–139 (2022). Publisher: Elsevier.

[5] Amodio, M. & Krishnaswamy, S. Dy, J. & Krause, A. (eds) Proceedings of the 35th International Conference on Machine Learning. MAGAN: Aligning biological manifolds, Vol. 80 of *Proceedings of Machine Learning Research*, 215–223 (PMLR, 2018).

[6] Wang, X., Wu, X., Hong, N. & Jin, W. Progress in single-cell multimodal sequencing and multi-omics data integration. Biophysical Reviews 16, 13–28 (2024).

[7] Wang, X. et al. BREM-SC: a bayesian random effects mixture model for joint clustering single cell multi-omics data. Nucleic Acids Research 48, 5814–5824 (2020).

[8] Singh, R., Hie, B. L., Narayan, A. & Berger, B. Schema: metric learning enables interpretable synthesis of heterogeneous single-cell modalities. Genome Biology 22, 131 (2021).

[9] Argelaguet, R. et al. MOFA+: a statistical framework for comprehensive integration of multi-modal single-cell data. Genome Biology 21, 111 (2020).

[10]. Liu, J., et al. Jointly defining cell types from multiple single-cell datasets using LIGER. Nature Protocols 15, 3632–3662 (2020). Publisher: Nature Publishing Group.

[11] Huizing, G.-J., Deutschmann, I. M., Peyŕe, G. & Cantini, L. Paired single-cell multi-omics data integration with Mowgli. Nature Communications 14, 7711 (2023). Publisher: Nature Publishing Group.

[12] Dou, J. et al. Bi-order multimodal integration of single-cell data. Genome Biology 23, 112 (2022).

[13]. Gayoso, A., et al. Joint probabilistic modeling of single-cell multi-omic data with totalVI. Nature Methods 18, 272–282 (2021). Publisher: Nature Publishing Group.

[14] Zuo, C. & Chen, L. Deep-joint-learning analysis model of single cell transcriptome and open chromatin accessibility data. Briefings in Bioinformatics 22, bbaa287 (2021).

[15] Qattous, H. et al. PaCMAP-embedded convolutional neural network for multi-omics data integration. Heliyon 10, e23195 (2024).

[16] Wen, H., et al. *Proceedings of the 28th ACM SIGKDD Conference on Knowledge Discovery and Data Mining*. *Graph neural networks for multimodal single-cell data integration*, KDD ’22, 4153–4163. ss (Association for Computing Machinery, New York, NY, USA, 2022).

[17] Athaya, T., Ripan, R. C., Li, X. & Hu, H. Multimodal deep learning approaches for single-cell multi-omics data integration. Briefings in Bioinformatics 24, bbad313 (2023).

[18] Koca, M. B., Nourani, E., Abbasoglu, F., Karadeniz, I. & Sevilgen, F. E. Graph convolutional network based virus-human protein-protein interaction prediction for novel viruses. Computational Biology and Chemistry 101, 107755 (2022).

[19] Szklarczyk, D. et al. The STRING database in 2017: quality-controlled pro- tein–protein association networks, made broadly accessible. Nucleic Acids Research 45, D362–D368 (2017).

[20] Jardine, L. et al. Blood and immune development in human fetal bone marrow and Down syndrome. Nature 598, 327–331 (2021).

[21] John, C. R., Watson, D., Barnes, M. R., Pitzalis, C. & Lewis, M. J. Spectrum: fast density-aware spectral clustering for single and multi-omic data. Bioinformatics 36, 1159–1166 (2020).

[22] Lopez, R., Regier, J., Cole, M. B., Jordan, M. I. & Yosef, N. Deep generative modeling for single-cell transcriptomics. Nature Methods 15, 1053–1058 (2018). Publisher: Nature Publishing Group.

[23] Chen, H., Ryu, J., Vinyard, M. E., Lerer, A. & Pinello, L. SIMBA: single-cell embedding along with features. Nature Methods 21, 1003–1013 (2024). Publisher: Nature Publishing Group.

[24] Cui, A. et al. Single-cell atlas of the liver myeloid compartment before and after cure of chronic viral hepatitis. Journal of Hepatology 80, 251–267 (2024).

[25] Wang, S. et al. An atlas of immune cell exhaustion in HIV-infected individuals revealed by single-cell transcriptomics. Emerging Microbes & Infections 9, 2333– 2347 (2020).

[26] Phaahla, N. G. et al. Chronic HIV-1 Infection Alters the Cellular Distribution of Fc*γ*RIIIa and the Functional Consequence of the Fc*γ*RIIIa-F158V Variant. Frontiers in Immunology 10, 735 (2019).

[27]. Poonia, B., Kijak, G. H. & Pauza, C. D. High Affinity Allele for the Gene of FCGR3A Is Risk Factor for HIV Infection and Progression. PLOS ONE 5, e15562 (2010). Publisher: Public Library of Science.

[28] Chen, Y. et al. An atlas of immune cell transcriptomes in human immunodeficiency virus-infected immunological non-responders identified marker genes that control viral replication. Chinese Medical Journal 136, 2694–2705 (2023).

[29] Ester, M., Kriegel, H.-P., Sander, J. & Xu, X. Proceedings of the Second International Conference on Knowledge Discovery and Data Mining. A density-based algorithm for discovering clusters in large spatial databases with noise, KDD’96, 226–231 (AAAI Press, 1996).

[30] Uhĺen, M., et al. The human secretome. Science Signaling 12, eaaz0274 (2019).

[31] Anlauf, M. et al. Vesicular monoamine transporter 2 (VMAT2) expression in hematopoietic cells and in patients with systemic mastocytosis. The Journal of Histochemistry and Cytochemistry: Official Journal of the Histochemistry Society 54, 201–213 (2006).

[32] Arock, M., Schneider, E., Boissan, M., Tricottet, V. & Dy, M. Differentiation of human basophils: an overview of recent advances and pending questions. Journal of Leukocyte Biology 71, 557–564 (2002).

[33] Welner, R. S. et al. Asynchronous RAG-1 expression during B lymphopoiesis. *Journal of immunology (Baltimore*, Md*. :* 1950) 183, 7768 (2009).

[34] Luger, D., et al. Expression of the B-Cell Receptor Component CD79a on Immature Myeloid Cells Contributes to Their Tumor Promoting Effects. PLOS ONE 8, e76115 (2013). Publisher: Public Library of Science.

[35] Kaiser, F. M. P. et al. IL-7 receptor signaling drives human B-cell progenitor differentiation and expansion. Blood 142, 1113–1130 (2023).

[36] Hoebeke, I. et al. T-, B- and NK-lymphoid, but not myeloid cells arise from human CD34(+)CD38(-)CD7(+) common lymphoid progenitors expressing lymphoid-specific genes. Leukemia 21, 311–319 (2007).

[37] Liang, K. L., Laurenti, E. & Taghon, T. Circulating IRF8-expressing CD123+CD127+ lymphoid progenitors: key players in human hematopoiesis. Trends in Immunology 44, 678–692 (2023).

[38] Stuart, T. et al. Comprehensive Integration of Single-Cell Data. Cell 177, 1888– 1902.e21 (2019).

[39] Guo, H., Barberi, T., Suresh, R. & Friedman, A. D. Progression from the Common Lymphoid Progenitor to B/Myeloid PreproB and ProB Precursors during B Lymphopoiesis Requires C/EBP*α*. *Journal of Immunology (Baltimore*, Md.: 1950) 201, 1692–1704 (2018).

[40] Morgan, D. & Tergaonkar, V. Unraveling B cell trajectories at single cell resolution. Trends in Immunology 43, 210–229 (2022).

[41]. Regev, A., et al. The Human Cell Atlas. eLife 6, e27041 (2017). Publisher: eLife Sciences Publications, Ltd.

[42] Bredikhin, D., Kats, I. & Stegle, O. MUON: multimodal omics analysis framework. Genome Biology 23, 42 (2022).

[43] Milacic, M. et al. The Reactome Pathway Knowledgebase 2024. Nucleic Acids Research 52, D672 (2023).

[44] Kullback, S. & Leibler, R. A. On Information and Sufficiency. The Annals of Mathematical Statistics 22, 79–86 (1951). Publisher: Institute of Mathematical Statistics.

[45] Wolf, F. A., Angerer, P. & Theis, F. J. SCANPY: large-scale single-cell gene expression data analysis. Genome Biology 19, 15 (2018).

[46] Hamilton, W. L., Ying, R. & Leskovec, J. *Proceedings of the 31st International Conference on Neural Information Processing Systems*. *Inductive representation learning on large graphs*, NIPS’17, 1025–1035 (Curran Associates Inc., Red Hook, NY, USA, 2017).

[47] Xiao, C., Chen, Y., Meng, Q., Wei, L. & Zhang, X. Benchmarking multi-omics integration algorithms across single-cell RNA and ATAC data. Briefings in Bioinformatics 25, bbae095 (2024).

[48] Luecken, M. D. et al. Benchmarking atlas-level data integration in single-cell genomics. Nature Methods 19, 41–50 (2022).

[49] Pedregosa, F. et al. Scikit-learn: Machine Learning in Python. J. Mach. Learn. Res. 12, 2825–2830 (2011).

[50]. Traag, V., Waltman, L. & Eck, N. J. v. From Louvain to Leiden: guaranteeing well-connected communities (2019). ArXiv:1810.08473.

[51] Kolberg, L. et al. g:Profiler-interoperable web service for functional enrichment analysis and gene identifier mapping (2023 update). Nucleic Acids Research 51, W207–W212 (2023).

[52]. Cui, T., et al. Gene–gene interaction detection with deep learning. Communications Biology 5, 1–12 (2022). Publisher: Nature Publishing Group.

[53] Barrett, T. et al. Ncbi geo: archive for functional genomics data sets—update. Nucleic Acids Research 41, D991–D995 (2012).

